# Metabolic Model-Based Analysis of the Emergence of Bacterial Cross-Feeding through Extensive Gene Loss

**DOI:** 10.1101/180208

**Authors:** Colin P McNally, Elhanan Borenstein

## Abstract

Metabolic dependencies between microbial species are common and have a significant impact on the assembly and resilience of microbial communities. However, the origins of such metabolic dependencies, the evolutionary forces that drive metabolic cross-feeding, and the impact of metabolic and genomic architecture on their emergence are not clear. To address these questions, we developed a novel simulation-based framework coupling a model of reductive evolution with a multi-species genome-scale model of microbial metabolism. We used this framework to model the evolution of a two-species microbial community, simulating thousands of independent evolutionary trajectories and investigating the link between genome reductive evolution and the emergence of metabolic interactions. Surprisingly, even though our model does not impose explicit selection for cooperation, metabolic dependencies emerged in nearly half of all evolutionary runs. Evolved dependencies involved cross-feeding of a diverse set of metabolites at varying frequencies, reflecting various constraints imposed by the metabolic network architecture. We additionally found metabolic ‘missed opportunities’, wherein species failed to capitalize on metabolites made available by their partners. When cross-feeding did evolve, it generally emerged immediately after a metabolite became available, but a complete dependence on such cross-fed metabolites often evolved relatively slowly. Examining the genes deleted and retained in each evolutionary trajectory and the timing of gene deletion events along these trajectories, we were further able to identify both genome-wide properties and specific gene retentions that were associated with metabolic phenotypes. Our findings provide insight into the evolution of cooperative metabolic interaction among microbial species, offering a unique view into the way such relationships could emerge in natural settings.

## Introduction

Most microorganisms in nature do not live in isolation but are rather part of complex communities [1]. Such communities inhabit a wide range of environments and have attracted significant interest in recent years due to an increased appreciation of their role in human health and environmental stewardship as well as their potential applications in industry, agriculture, and medicine [2–5]. The various microbial species that form these communities not only share a common environment, but rather interact with other community members in various ways including competition for extracellular nutrients, cooperation through metabolite cross-feeding, signaling, biofilm formation, and antimicrobial secretion [1,6]. Such interactions allow community members to impact each other’s behavior and play an important role in shaping community structure and activity. A better understanding of how these interactions emerge through ecological and evolutionary dynamics, how they are maintained or lost, and how they impact community-level behavior is therefore crucial for both understanding the forces that have shaped current natural communities and for designing synthetic communities or targeted modulation of natural communities.

Perhaps the most intriguing form of microbial interaction is inter-species cooperation. The prevalence of cooperative interaction is evident from the large number of microbes that cannot be individually cultured, suggesting that they are reliant on symbiotic interactions with other members of their communities [7]. In the context of metabolism, cooperation often takes the form of cross-feeding, where one species secretes metabolites that other species uptake and utilize. Indeed, metabolic cross-feeding has been found to occur in a wide variety of environments and between diverse species [8], often benefiting both partners [9]. For example, *Bifidobacterium* species in the gut microbiota regularly cross-feed fermentation products and partial digestion byproducts of polysaccharides to butyrate-forming bacteria [10,11]. Furthermore, evidence suggests that metabolic cooperation drives species co-occurrence in diverse microbial communities [12].

Notably, however, metabolic cooperation is not limited to complex communities and has also been demonstrated in small two- or three-species communities, such as those occupying various insect hosts. For example, it was shown that the two endosymbionts that occupy a sharpshooter insect (and the insect host) each lack necessary steps of several amino acid synthesis pathways, and consequently only when the three organisms grow together can they synthesize the entire complement of amino-acids [13,14]. Similarly, it was shown that two endosymbiotic bacteria that inhabit a *Cicadoidea* host had recently diverged into 3 species, with metabolic complementarity between the two recently split lineages [15]. Studying the facultative symbionts of another host, the sap-feeding whitefly, it was shown that pairs of symbionts with greater metabolic complementarity are more likely to co-occur in a host [16]. Such tightly coupled metabolic systems, where two or three species strongly depend on each other for survival, could be viewed as an idealized model of microbial cooperation and accordingly have been extensively studied. Similar tight cross-feeding behavior between a small number of species has been successfully engineered in laboratory conditions, for example allowing auxotrophic strains of *Escherichia coli* to survive in co-culture [17–19]. Cross-feeding has also been engineered between different strains of a yeast species [20], and even between species as divergent as yeast and algae [21].

This prevalence of cooperation is, however, somewhat surprising given a large body of work suggesting that cooperation should be challenging to evolve [22,23]. Specifically, the emergence of cheaters or the potential loss of species that provide essential metabolites could hinder the evolution of cooperative interaction. One possible explanation for the emergence of cooperation in the face of these challenges is the Black Queen hypothesis [24], positing that symbiotic interactions through gene loss could evolve when a function is biosynthetically costly but has a leaky benefit [25]. This mechanism, however, may not be applicable in fixed communities with small population size, where selection is relatively weak [26]. In such communities (such as insect endosymbionts), cooperation likely emerges not through selection by rather by chance as a consequence of extreme genome reduction [27]. Indeed, most insect endosymbionts have extremely small genomes (some of which are the smallest bacterial genomes known) that are majorly reduced compared to their closest free-living relatives [28].

Importantly, however, despite the prevalence and diversity of such obligate endosymbionts, the process and determinants through which extensive long term genome reduction gives rise to metabolic cross-feeding are not clear and their study is challenging. Experimental evolution studies have aimed to examine the evolution of metabolic cooperation, evolving a population seeded with a single species and demonstrating the emergence of cooperative interactions between divergent polymorphic subsets of the populations [29,30]. Similar experimental evolution studies that have focused on populations seeded with multiple species that were initially cooperative have also shown that such species evolve to be both more efficient at and more dependent on that cooperation [31,32]. However, such evolutionary experiments are limited in both duration of evolution (which precludes studying extreme long term genomic reduction) and number of replicates (which hinders development of a more comprehensive view of microbial evolution and identification of general principles in the emergence of species interaction). Computational methods, on the other hand, allow long time scales to be easily modeled and thus may be more applicable. For example, prior studies determined the dispersal properties that favor cross-feeding by modeling the co-evolution of microbial strains where the various genotypes determined which compounds were secreted to a shared environment [33]. Another study used a mathematical model of two species that can increase each other’s fitness, aiming to identify the processes underlying the evolution of cooperation [22]. Such models have also been used to study the impact of genetic variability on synergistic effects between partner species [34], and to investigate the effect of population density on the emergence of cross-feeding [35]. These studies have produced useful insights, but tend to explicitly model interaction in a non-mechanistic way and thus are limited in utility for studying how a genomic evolutionary process (e.g., genome reduction through genetic drift) could lead to the evolution of cross-feeding.

To address this gap, in this study, we utilized a model of microbial evolution over a *long time scale* coupled with a *mechanistic model* of multi-species microbial growth. Our model is inspired by a previous computational study that modeled reductive evolution of a single endosymbiont species [36]. The authors investigated how historical contingency between gene deletions affects future genome reduction and the nature of the minimal genomes that result. This framework models metabolism on a genome scale and is able to simulate long evolutionary time scales, but is also amenable to highly replicate simulations to infer general principles. In the work we present here, we extended this evolutionary framework to a co-culture model of two species, using a mechanistic model of microbial growth in co-culture which is based on a recently introduced multi-species genome-scale metabolic modeling approach [37].

We specifically aim to examine whether species interaction can emerge without explicit selection, and which mechanisms can drive a selfishly evolving species to support a dependent species. To this end, we model the evolution of cross-feeding in a simple multi-species community. Notably, we do not model the process by which an evolving population bifurcates into multiple subpopulations, but rather explicitly assume the community harbors two interacting but evolutionary isolated species (e.g., following an initial split), each undergoing an extreme reductive evolution process. We further assume that these two species co-exist over a long time scale, without one outcompeting the other. Using genome scale metabolic models to model each of these evolving species, we were able to to directly investigate how the architecture of the metabolic and genetic network affects the evolution of cross-feeding interactions, and how evolved genomes reflect the type of interaction that emerged. Using this framework, we simulated thousands of independent evolutionary trajectories, tracked the emergence of metabolic cross-feeding, and carefully analyzed the evolving species. We aimed to address several fundamental questions concerning the evolution of cooperation under such an evolutionary regime. Can species interaction emerge without explicit selection for it? How do species evolve to depend on available metabolites and how rapidly will such dependencies emerge? What are the mechanisms that drive a selfishly evolving species to support a dependent species? How does the architecture of the metabolic and genetic networks affect the evolution of such interactions, and how do the evolved genomes reflect the type of interaction that emerged? And finally, how does evolution in a community shape co-evolutionary dynamics? Addressing these questions could have profound impact both on our understanding of the evolution of species interactions and on our ability to design and construct stable microbial communities for medical, agricultural, and industrial application.

## Results

### A framework for modeling the evolution of species interactions

To study the emergence of metabolic species interaction in bacteria we developed a computational framework that integrates models of microbial co-evolution, metabolic activity, and ecological interaction (Fig 1). Briefly, in this framework, we model a community comprised of two generalist species growing in a shared environment (and that can therefore exchange metabolites) that go through a reductive evolution process. Our reductive evolution model is inspired by a previously introduced model of reductive evolution of a single endosymbiont species [36]. In our model evolution is an iterative process in which a gene is first chosen at random from either of the two species for deletion (Fig 1A). The fitness effect of losing that gene (in the context of the community) is calculated using a co-culture metabolic model (described below). If the decrease in fitness to the species losing this gene does not exceed a predefined threshold the deletion is assumed to fix; otherwise the deletion is assumed to be selected against and is reverted. Importantly, during the course of this co-evolutionary process, the presence of each of the two species in the community can markedly impact the evolution of the other (and specially, the set of genes that can be deleted). This process repeats until no more genes can be deleted from either species.

**Figure 1:**
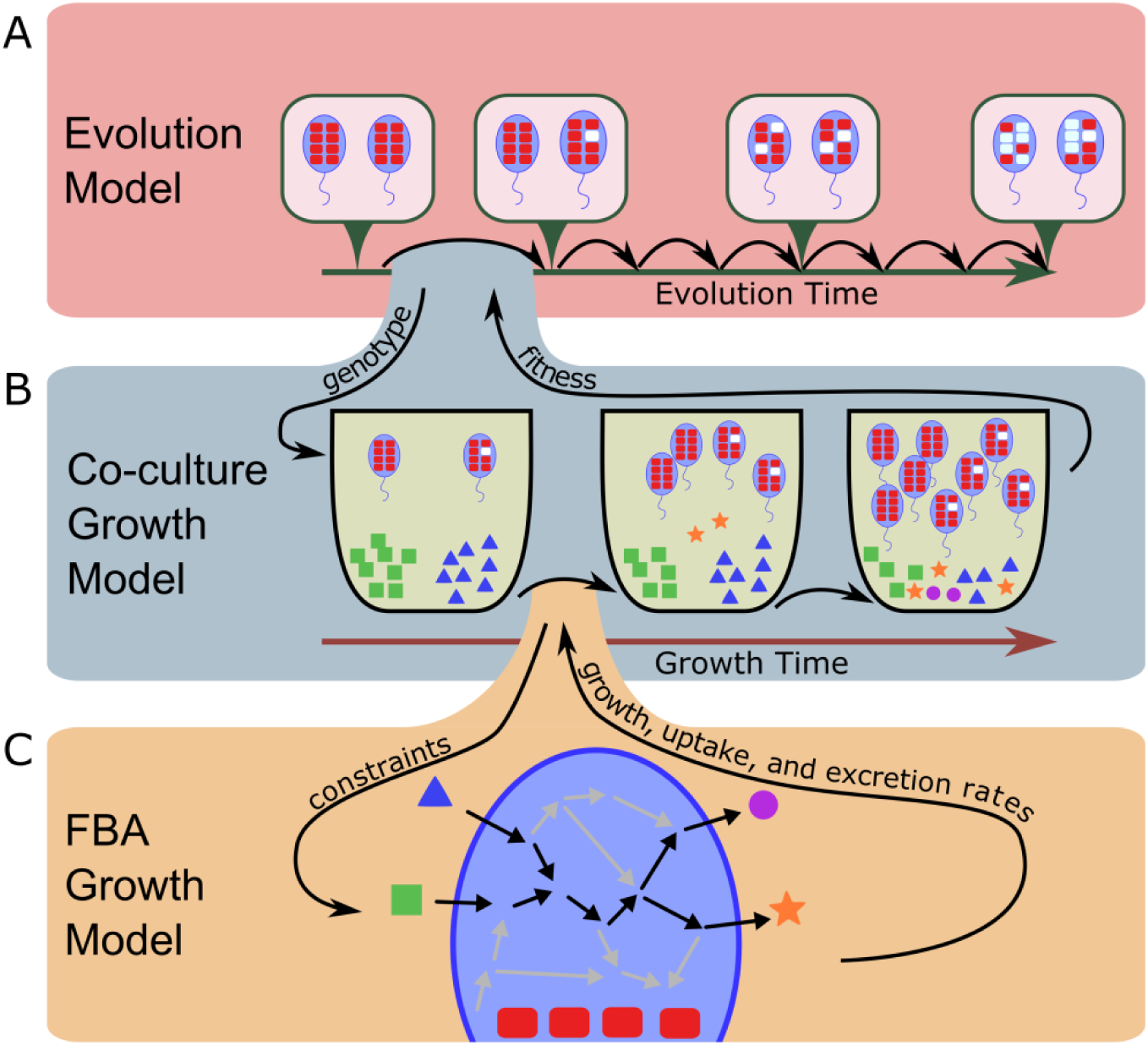
A framework for modeling the evolution of species interaction. **(A)** To model reductive evolution, genes are iteratively chosen at random as candidate for deletion, the fitness effect of their deletion is evaluated (using a co-culture growth model; see panel B), and if the fitness effect is relatively small, these genes are deleted. **(B)** The co-culture growth model simulates the growth of the two species in a well-mixed shared environment, and is based on a previously introduced dynamic multi-species model [37]. This model iteratively infers the behavior of each species in the shared environment based on an FBA approach (see panel C). The predicted growth of each species and the predicted rates at which it uptakes and excretes various metabolite are used to update the abundances of species in the co-culture and the concentration of metabolites in the shared environment over time. **(C)** An FBA model is used to predict the growth of each species in a given environment based on the set of metabolic reactions and constrains encoded by the species and the concentration of metabolites in its environment.

Specifically, to model growth in co-culture and to determine the fitness consequence of gene deletions while accounting for the way the presence of one species in the community may impact the fitness of the other, we used a previously introduced co-culture metabolic modeling framework [37]. This framework employs dynamic Flux Balance Analysis (FBA) to predict the metabolic activity and growth of two species in a shared environment over discrete time points (Fig 1B). Briefly, at each time point and for each species in the co-culture, this framework uses a genome-scale metabolic model of the species (based on its metabolic capacity as determined by the set of genes present in its genome), the current concentration of metabolites in the environment, and a Flux Balance Analysis to predict the species’ behavior, including its growth rate and the rate at which it imports and excretes various metabolites [38] (Fig 1C). The estimated growth rates of the two species are then used to update the abundances of the species in the community and the predicted uptake and excretion fluxes are used to update the concentration of metabolites in the shared environment. Growth is simulated over several time points and the growth rates of each species at the last time point are used as proxies for their fitness.

To classify the interaction between the two species in each community and at each evolutionary step, we also simulated and evaluated the growth of each of the two species in isolation (i.e., in mono-culture). We define a species as being dependent on its partner if it can grow in co-culture but not in mono-culture (zero fitness). Accordingly, we distinguish between three possible types of interaction a given community can exhibit: (i) *Independent* – neither species is dependent on the other, (ii) *commensal* – one species (*‘dependent’*) is dependent on the other species but the other (*‘provider’*) is not, and (iii) *mutualistic* – both species are dependent on each other. The observed interaction type at the end of the simulation run (i.e., when both species reach minimal genomes) was used to label each evolutionary trajectory (Fig 2A). Notably, in some cases, one of the two species can go through a catastrophic drop of fitness (>50%) even in co-culture (e.g., due to a change in the *other* species’ behavior that limits the availability of a metabolite it requires). In such cases, that species was considered to have gone extinct and the simulation was labeled as a collapsed community. A detailed description of the framework is provided in Methods.

**Figure 2:**
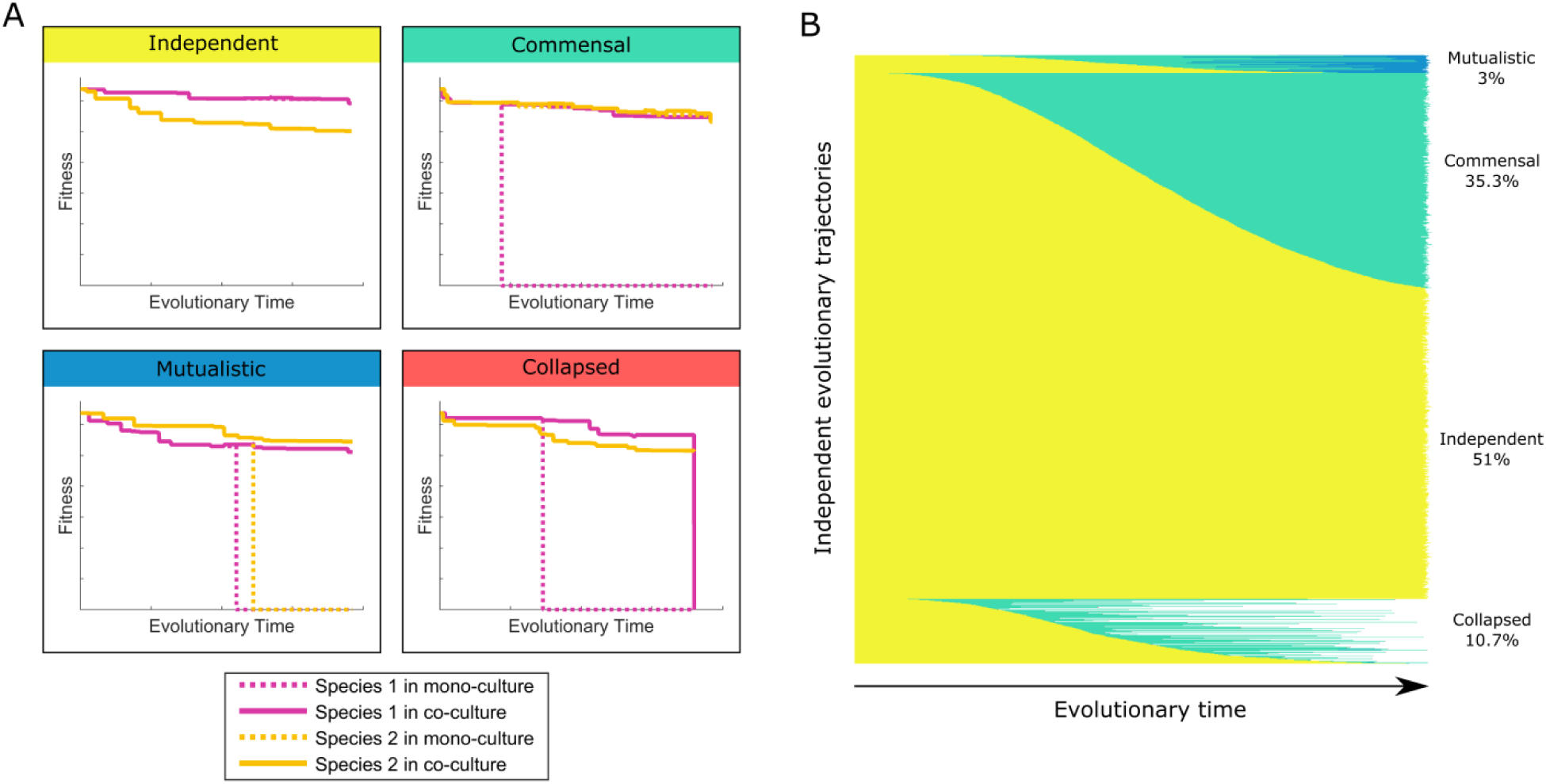
Types of interactions and their emergence over time. **(A)** Evolutionary simulations could result in one of four unique outcomes, determined by the ability of evolved species to grow in mono-culture and co-culture. Plotted are examples of each of these four outcomes, illustrating the fitness of each of the two species in mono-culture and in co-culture over evolutionary time. **(B)** The changes in interaction type over time for all 16,317 simulation runs. Each horizontal bar represents a single simulation run, and the color corresponds to the interaction type using the same colors as the titles in panel A.

### The emergence and prevalence of species interaction

We used the framework described above to simulate the evolution of a simple community comprising two species that go through a reductive evolution process. We assumed that the community composition is fixed as a two genotype community (i.e., with no new species migrating into the community and no standing genetic diversity). In the simulation below, we initialized the community with two identical *E. coli* strains (as a generalist model species). Such a scenario may represent the evolutionary trajectory of two obligate symbionts that may have diverged from a common ancestor.

We simulated 16,317 independent evolutionary trajectories (Methods) and for each simulation examined the evolved two-species community and the interaction between the two evolved species (Fig 2A). Surprisingly, although our framework does not impose an explicit pressure toward species interaction, we found that a substantial fraction of simulations resulted in a community with some sort of metabolic dependency between the two species. Specifically, 35% of the simulations ended with a commensal community, and 3.2% of the simulations ended with a mutualistic community (Fig 2B). In 10.7% of the simulation, the community collapsed as described above. The remaining 51.1% of simulations ended with independent communities in which both species were still capable of independent growth. Using different fitness cutoffs for allowing deleterious gene deletions to fix affected the ratio of the different interaction types, with a more stringent cutoff resulting in more independent communities and a less stringent cutoff resulting in more commensal and mutualistic communities (see Supporting Text). In natural communities the strength of selection against deleterious gene deletion reflects multiple factors, ranging from population size to environment stability, which therefore indirectly affects the likelihood of emergent cooperation.

### An example of an emergent cross-feeding interaction

Next, we set out to examine the specific genes and metabolic processes involved in emergent interactions. Before exploring large-scale patterns concerning the mechanisms involved in species interaction, we set out to characterize in detail one evolved community as an example of the kind of metabolic interaction that could emerge and the gene deletions that underlie such an interaction. We specifically focused on one simulation run where the two evolved species (arbitrarily referred to below as species A and species B) exhibited a mutualistic interaction. In this simulation, species A had retained only 306 genes and species B had retained only 304 genes (compared to 1260 genes in the ancestor species) and neither could grow in mono-culture. The two species, however, could still grow in co-culture (albeit at only 78% and 73% of the ancestor’s growth rate, respectively).

To identify metabolites that may be involved in cross-feeding, we first detected all the metabolites found in the medium when the two evolved species grew in co-culture and that were not included in the initial growth medium. We then tried growing each of the two evolved species in mono-culture by augmenting the initial minimal growth medium with one or more of these candidate cross-feeding metabolites. We found that species A could grow on the initial medium once tyrosine was added (at 78% of the ancestor’s growth rate). Species B could similarly grow on the initial medium once thymidine was added (at 72% of the ancestor’s growth rate).

We further examined the fluxes through the metabolic models of the evolved species and compared them to the fluxes observed in the ancestor species, to identify the specific gene deletions that gave rise to these dependencies (Fig 3). We found that species A became dependent on external tyrosine due to a loss of the gene *tyrA*, which is necessary for tyrosine synthesis [39]. Indeed, species A’s loss of *tyrA* occurred at the exact same point in the evolutionary trajectory as its loss of the ability to grow in mono-culture. Similarly, Species B became dependent on external thymidine due to a loss of the gene *thyA*, which is necessary for dTMP synthesis [40]. Notably, we were also able to identify the evolved mechanisms that allowed each of the two species to excrete the metabolite necessary for growth of the other species. Specifically, species A started excreting thymidine due to a loss of the gene *cmk*, which is necessary to phosphorylate CMP to CDP [41]. The loss of several other reactions prevented species A from converting CMP to cytidine, uridine, uridine monophosphate, excreted uracil, or thymine, which resulted in species A only being able to eliminate excess CMP by converting it to thymidine and excreting it. Indeed, a *cmk* deletion in *E. coli* has been shown experimentally to result in 30-fold elevated CMP and dCMP pools relative to wild-type [41]. Species B similarly excreted tyrosine due to an overproduction of this metabolite following a complex combination of gene losses that resulted in elevated activation of the pentose phosphate pathway and converting excess erythrose-4-phosphate into tyrosine. This community provides examples of specific mechanisms that drive one species to support another via cross-feeding, and those that are involved in the second species becoming dependent on these cross-fed metabolites. We will next examine the prevalence of these and other specific metabolic interactions.

**Figure 3:**
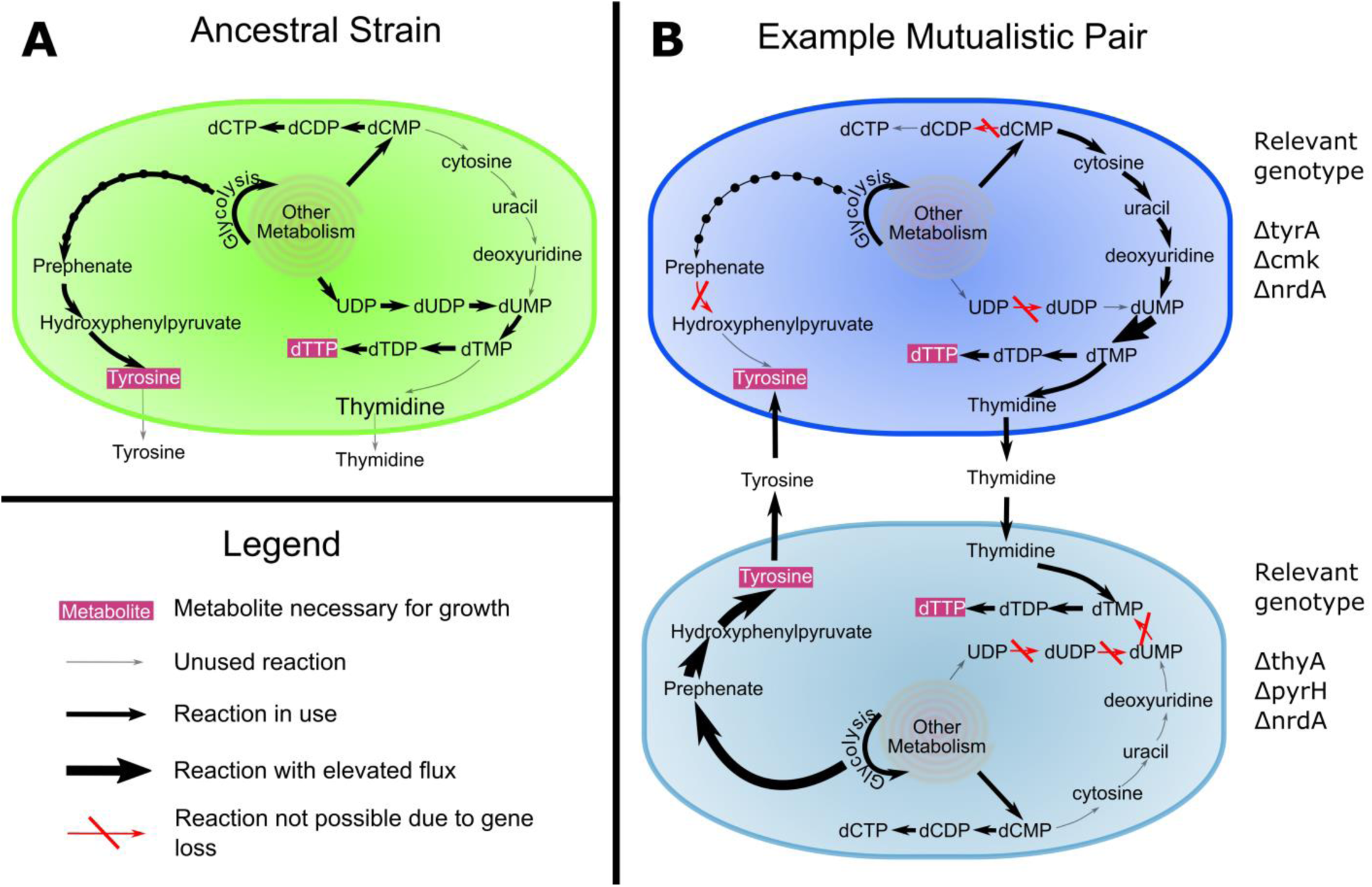
Example of an evolved mutualistic community. **(A)** In the ancestral species tyrosine is produced through the shikimate pathway and dTTP is produced from UDP. **(B)** In this example evolved mutualistic community, deletions in both species have led to obligate cross-feeding of tyrosine and thymidine. The relevant gene deletions and their impact on metabolic fluxes in each species are highlighted.

### Metabolite cross-feeding in evolved pairs

After characterizing one cooperating pair in detail, we moved on to examine the complete set of communities evolved by our model, focusing specifically on identifying the metabolites underlying emergent species interactions. To infer such cross-fed metabolites, we simulated the growth of each evolved dependent species on minimal media supplemented with various metabolites and determined the minimal set of supplemented metabolites required for growth (as described above; see Methods). This set was assumed to represent essential metabolites provided by the partner.

We found that the majority of dependent species (94.3%) required only a single essential metabolite to be cross fed from their partner, with only a small fraction of dependent species requiring two or three such metabolites (5.5% and 0.2% respectively), and no species requiring more than three. Formate was the most common essential metabolite (65% in commensal dependent species; see Fig 4A), followed by Tyrosine (17.6%) and Phenylalanine (6.4%). Notably, the dependence on a single (or very few) metabolites reported above contrasts observations made in several insect symbionts systems where cooperating symbionts exchange multiple essential compounds (and see Discussion below), yet the exchange of aromatic amino-acids is in agreement with cross-fed metabolites often observed in such systems [13,42].

**Figure 4:**
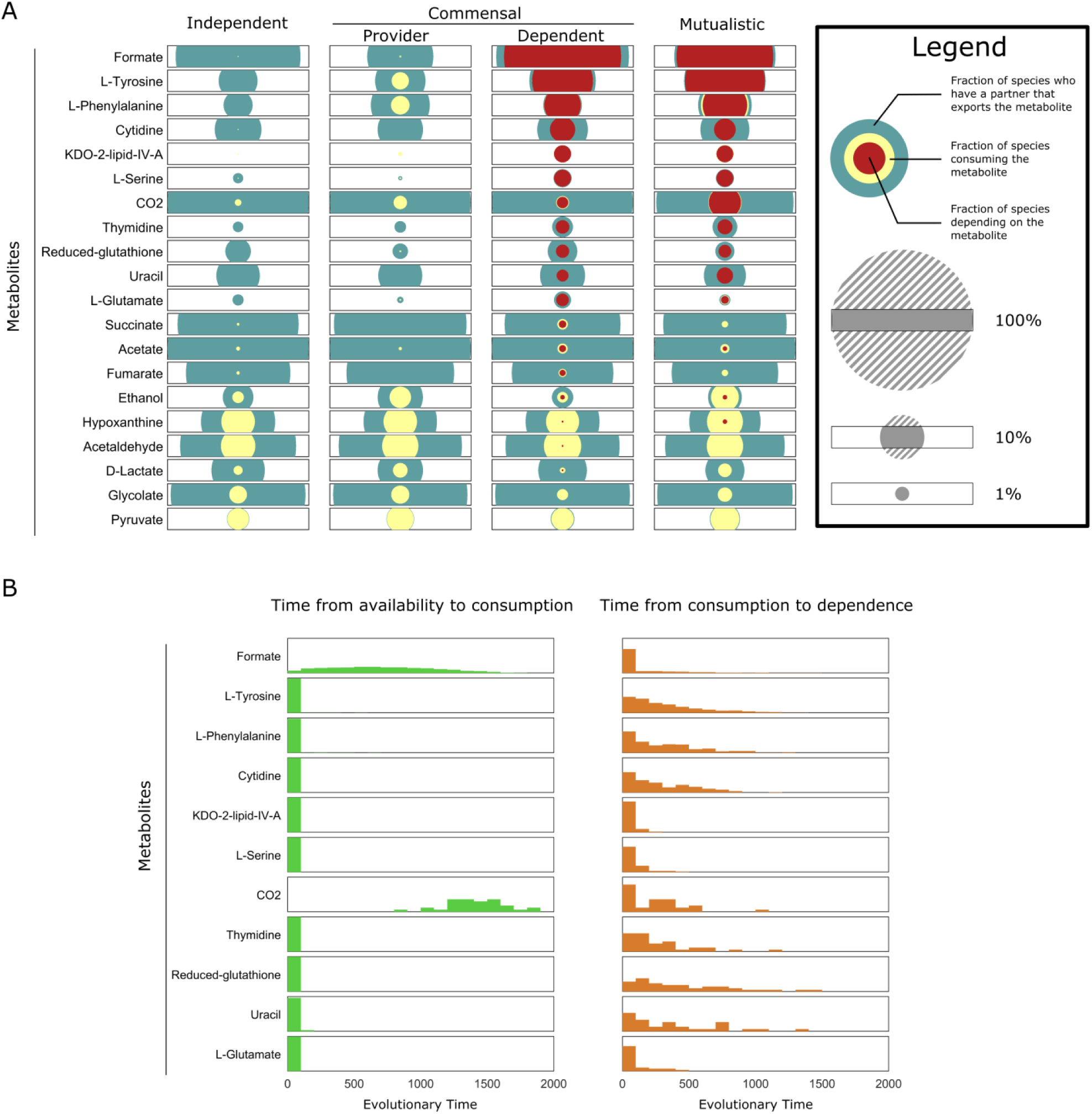
Frequencies of metabolites’ availability, cross-feeding, and dependence and the timing of their emergence. **(A)** The frequencies of metabolites’ availability, cross-feeding, and dependence are shown for species of each interaction type and for each metabolite. Each set of nested circles shows the frequency at which the given metabolite is produced by their partner species and hence available for uptake (blue), the frequency at which this metabolite is utilized by the species through cross-feeding (yellow), and the frequency at which this metabolites is depended on (red). The area of the circle scales with the frequency, but for visualization purposes the portions of the circle extending beyond the rectangular box are not shown. **(B)** The distributions of evolutionary time (measured as number of gene deletions) elapsed between the different stages of metabolic interaction, for commensal species dependent on a single metabolite.

Complete dependence on cross-fed metabolites (such as those identified above) is the most defining feature of species interactions in these communities, but may represent an extreme form of interaction. Clearly, cross-feeding can be beneficial to a species even when it is not essential for growth, and in fact this form of cross-feeding may be a common precursor state of complete dependence. To detect such non-essential cross-fed metabolites we examined transporter fluxes and identified metabolites that are being excreted by one species in the community and uptaked by the other (Fig 4A, yellow circles). Indeed, in addition to the essential metabolites identified above, our analysis revealed multiple metabolites that are being cross-fed but are non-essential to growth. Such non-essential cross-feeding often involved metabolites that were rarely if ever depended on (such as acetaldehyde and pyruvate) and were observed at similar frequencies in species across all interaction types. Interestingly, however, we also detected non-essential cross-feeding of metabolites that were commonly depended upon, but these were rare in independent communities and occurred surprisingly often in commensal communities where *providers* were cross-fed such metabolites by the dependent partner (see for example, tyrosine and phenylalanine in Fig 4A). This finding suggests that species cooperation may involve two species that evolve a similar metabolic strategy (and therefore have the potential to both excrete and utilize a similar set of metabolites). In such cases, cross-feeding is likely to emerge, first as a non-essential process, which may later evolve into species commensal or mutual dependence. To confirm this hypothesis, we specifically examined, for each dependent species, the time that elapsed from when this species started consuming a metabolite via cross-feeding to when it became dependent on that metabolite. We find that indeed, in most cases dependence does not immediately follow cross-feeding, and there is often a substantial delay between cross-feeding and dependency (Fig 4B).

Clearly, uptaking a metabolite is only possible if the partner species is producing that metabolite and excreting it to the shared environment, thereby providing an opportunity for cross-feeding. We additionally quantified the frequency and time at which such opportunities arose, regardless of whether the metabolite was consumed or not (Fig 4A, blue circles). We found that metabolites vary greatly in the frequency at which they are excreted, and in a way that is not fully correlated with the frequency at which they are cross-fed or dependent on. For example, various metabolites, including cytidine, succinate, and acetate, are excreted at relatively similar frequencies in all interaction types, suggesting that dependency on these metabolites is not limited by their availability. Conversely, other metabolites, such as serine and thymidine are rarely excreted in independent communities, suggesting that the availability of these metabolites often lead to cross-feeding and dependency on them. Most importantly, while cross feeding often started almost immediately after the metabolite was available (in cases in which it occurred; see Fig 4B), in many cases evolving species failed to utilize available metabolites, thus completely missing cross-feeding (both essential and non-essential) opportunities (Fig 4A). This finding implies an intriguing dichotomy where available opportunities are either utilized immediately or are not utilized at all, potentially due to evolutionary constraints.

We finally examined the total number of different metabolites being excreted by each species over time, hypothesizing that species that excrete useful metabolites early on are more likely to become provider species. Surprisingly, however, we found that during the first half of the evolutionary process future providers in fact tend to excrete a similar or even a smaller number of metabolites on average compared to future dependents (Fig S1), and only toward the end of the evolutionary process did providers excrete more metabolites than dependent species. This pattern could suggest that species that eventually became dependent were less optimal early on, excreting more waste products, and that this wasteful behavior may have led to the development of dependence. Notably, all species gradually excrete more metabolites over the course of the evolution process, likely reflecting more complex growth strategies imposed by their shrinking genome).

### The genomic basis of evolved interactions

Our mechanistic model of microbial metabolism allows us to move beyond a phenotype-level description of evolved communities and to directly investigate patterns of genome evolution and identify genomic mechanisms involved in species interactions. We first examined the number of genes that were retained or lost in different simulations to explore the relationship between genome size (in terms of the number of genes retained) and species interaction. Surprisingly, with the exception of collapsed communities, evolutionary trajectories exhibited a markedly low variation in the total number of genes retained (note, for example, the similar length of the simulation runs illustrated in Fig 2B), with an average of 298.6±4.4 genes retained in each species. Yet, when comparing species from communities of different interaction types, we found that both dependent and mutualistic species had slightly but significantly smaller genomes compared to independent species (*P* < 10^-30^ and *P* < 10^-9^ respectively; two sample t-test; Fig 5A). Moreover, within commensal communities, the genomes of dependent species were slightly but significantly smaller than the genomes of provider species (*P* < 10^-30^). Notably, while these differences in *average* genome size were generally very small (often less than a single gene), the differences between the *smallest* genomes observed in the dependent or mutualistic species and the smallest genomes observed in provider or independent species was much larger (278-287 genes; Fig 5A). These results are consistent with the idea that cross-feeding allows dependent species to lose genes they would not be able to lose otherwise [14,43]. Moreover, provider species had on average a slightly larger genomes than independent species (*P* < 10^-3^), suggesting that provider species are a potential consequence of evolutionary trajectories that ended with larger minimal genomes. Notably, examining the genes retained across the evolved species, we found that of the initial 1260 genes present in the ancestral species, 560 were always lost and 149 were always retained, with only 551 genes being retained at intermediate frequencies (Fig 5B). The specific subset of these 551 genes that were retained in each evolved species therefore determines the types of interaction that emerged, and indeed a statistical analysis was able to detect specific genes whose retention or loss was associated with specific types of interaction (see Supporting Text).

**Figure 5:**
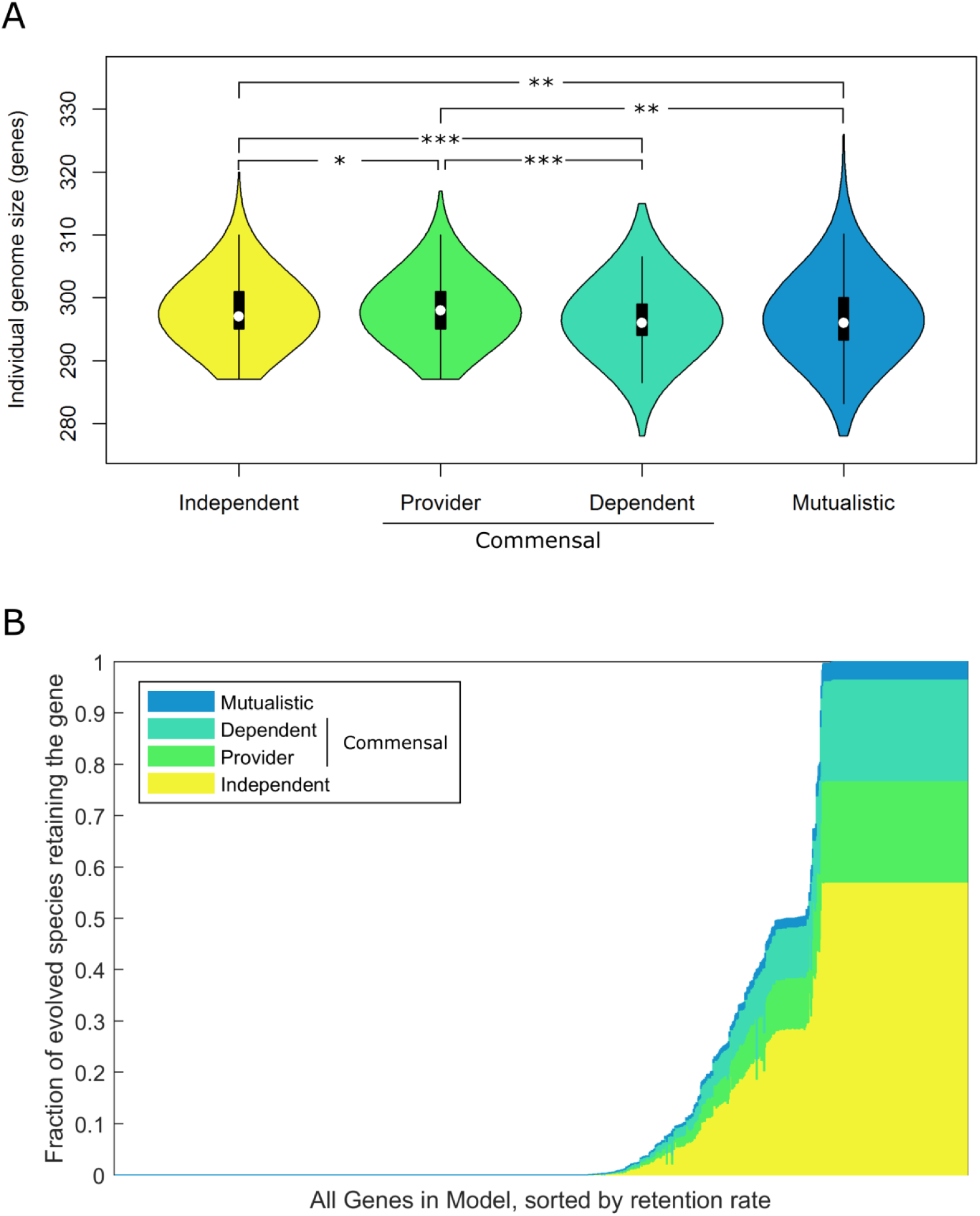
Genome size and gene retention frequency in evolved genomes. **(A)** Distributions of the genome size of evolved species from each interaction type. (*: *P* < 10^-3^ ; **: *P* < 10^-9^ ; ***: *P* < 10^-30^) **(B)** The distribution of retention rates of different genes in the model. Each gene is plotted as a vertical bar with height equal to the fraction of species in non-collapsed simulations that retained it. Each bar is colored by the fraction of species retaining that gene that are in each of the different interaction types.

We next examined how similar, on average, are the sets of genes retained between the two partners in each community. We found that mutualistic species were less similar to each other than independent species (*P* < 10^-20^; two sample t-test). This finding suggests that the evolution of metabolic dependency is associated with a process of functional diversification, where each of the two species retains certain metabolic capacities that the other species has lost. To further investigate this diversification process we turned our attention again to commensal communities, in which the two species can be labeled clearly as dependent and provider and therefore the direction of dependency is clear. Such communities also represent an intermediate level of interaction as compared to mutualistic communities, in which each species is acting as both provider and dependent, which complicates dissection of the mechanism of interaction. Indeed, the two partner species in commensal communities were more similar to one another than species in mutualistic communities (*P* = 1.4 x 10^-7^) but more divergent than species in independent communities (*P* < 10^-20^).

To better understand the diversification between providers and dependent species in commensal communities, we compared the set of genes retained in providers vs. those retained in dependent, identifying 80 genes that are more frequently retained in the provider and 41 that are more frequently retained in the dependent species. For example, *pflA*, a pyruvate formate lyase, was retained in 55.5% of providers but only 16.8% of dependents, whereas *aceE* and *aceF*, both components of the pyruvate dehydrogenase complex, were retained in 81.1% in dependent species and only 19.0% in provider species (these genes were also those with the greatest differential retention rate). This previous analysis identified genes more often retained in the provider or dependent species, yet ignored information about which of these retentions co-occurred in the same provider-dependent community. Applying a hypergeometric test (see Methods), we therefore identified a set of 263 pairs of genes that are significantly exclusively-retained in commensal communities (i.e., the dependent species retained the first gene when the provider lost the second gene or vice-versa more often than expected by chance; Fig S2). We further found that this set of exclusively-retained gene pairs was enriched for pairs that shared a pathway annotation (P < 10^-4^; permutation-based test), suggesting complementation at the pathway level.

We finally set out to examine the dynamics of gene deletion events in commensal communities, specifically focusing on the *order* in which deletions occurred in the provider and dependent species and aiming to detect dependencies between these deletion events that could highlight key evolutionary steps on the route to cross-feeding. To this end, we used a permutation-based analysis to identify instances where a gene in one species tended to be deleted after another gene was deleted in the partner species (see Methods). We identified 9 such gene pairs (at 1% FDR), all of which involved a gene being deleted in the dependent species significantly more often after a different gene was first deleted in the provider. Specifically, deletion of the *tyrA* gene in the dependent often followed deletion of a set of genes (*talA, talB, aroP*, and *pheP*) in the provider. *talA* and *talB* catalyze a reaction connecting glycolysis to the pentose phosphate pathway, and their deletion likely disrupts central carbon metabolism and diverts excess flux toward aromatic amino acid biosynthesis. Similarly, *aroP* and *pheP* are both transporters capable of transporting phenylalanine, and their deletion potentially prompts the excretion of tyrosine instead of phenylalanine. These deletions therefore promote over production and excretion of tyrosine by the provider, allowing the dependent to lose *tyrA,* a gene necessary for tyrosine synthesis. The deletion of the *pheA,* a gene necessary for phenylalanine synthesis in the dependent, was also found to follow the deletion of *talA* and *talB* in the provider, which is not surprising given the similarity in the biosynthesis pathways of these two amino acids. Finally the deletion of *pyrG* in the dependent tended to follow the deletion of *cdd, cmk,* and *codA* in the provider. The deletion of *cmk* (necessary for recycling CMP into CTP) and of *cdd* and *codA* (catalyzing reactions that could convert CMP or related products into other bases) could result in cytidine excretion and accordingly allows the dependent to lose *pyrG* (a component of CTP synthase) which creates a dependency on cytidine (Fig S3). To further examine the mechanism involved in these interactions, we tested whether the deletions of these key genes are sufficient to cause over-production and excretion of the relevant metabolites. Indeed, we found that deletion of *cdd*, *cmk*, and *codA* in the ancestral species (i.e., without any additional gene deletions) led to cytidine excretion. Deletion of *talA, talB, aroP, and pheP* in the ancestral species, however, was insufficient to cause excretion of either phenylalanine or tyrosine, suggesting that additional gene deletions are necessary to give rise to this phenotype.

### Linking genome evolution to metabolite cross-feeding

Having identified both the metabolites involved in cross-feeding and the genes involved in the emergence of species interaction, we finally turned to examine the association between specific gene retention or loss events and the cross-feeding of specific metabolites. Towards this end we again considered the set of all commensal communities and, for each of the 14 cross-fed metabolites that were depended upon at least 10 times, identified genes whose retention or deletion are significantly correlated with the excretion of this metabolite in the provider or with the dependency on this metabolite in the dependent (see methods). In total we identified 384 gene-metabolite associations in provider species, including 226 significant gene retentions and 158 significant gene deletions associated with essential metabolite excretion (Fig 6; χ^2^ test, 1% FDR). We similarly identified 459 gene-metabolite associations in dependent species (277 retentions and 182 deletions) associated with metabolite dependency. In total, retention or loss of 197 of the 510 variable genes was significantly associated with excretion of or dependence on at least one metabolite. Moreover, many genes were associated with the excretion of or dependence on more than one metabolite, likely reflecting the interconnectedness of the metabolic network where the loss of a key gene could affect multiple metabolic phenotypes.

**Figure 6:**
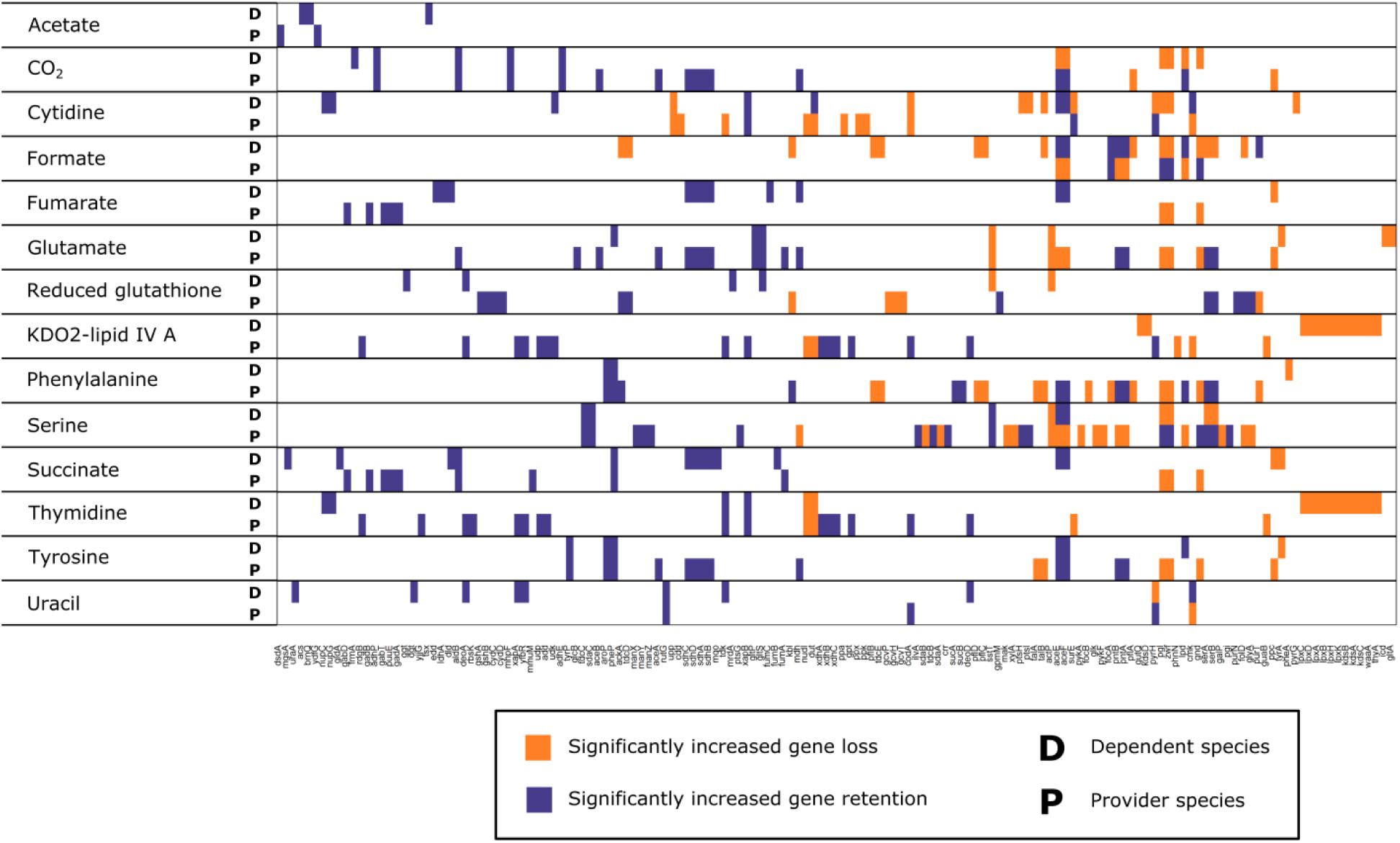
Associations between providing or depending on specific metabolites and deletion or retention of specific genes. Orange (purple) bars indicate genes that are lost (retained) significantly more often in species that provide or depend on a specific metabolites compared to independent species. Only associations with an absolute difference in gene retention frequency between the provider/dependent species and independent species ≥10% are shown. Genes are sorted by deletion frequency among all species from non-collapsed communities.

Interestingly, certain genes were associated with both the excretion of a metabolite by the provider and with the dependence on that same metabolite in the dependent species. For example, retention of *tyrP* and *aroP* (aromatic amino acid transporters) were associated with both providing tyrosine and dependence on tyrosine, as cross-feeding of that metabolite requires that both species in the pair could exchange it with the shared environment. In other cases the same gene was associated with a metabolite being excreted by the provider and utilized by the dependent but in a different direction. For example *serA* and *serB* – genes involved in serine biosynthesis – tended to be lost in species dependent on serine, but retained in species producing it. In rare cases there were genes whose loss was associated with both dependence and providing of a specific metabolite. For example, the loss of the gene *codA* was associated with both excretion of cytidine by providers and dependency on cytidine in dependent species (and see our analysis of that gene above). Combined, these associations suggest a complex link between evolutionary gene loss events and the emergence of metabolic species interactions and highlight the multitude of ways through which such interactions could evolve.

## Discussion

In this study we investigated the potential for metabolic mutualism to emerge between species inhabiting a shared, isolated environment and undergoing continual gene loss. We found that cross-feeding interactions emerged frequently (although two-way mutualistic interactions were much rarer). By examining associations between the loss and retention of genes and cross-fed metabolites we were able to elucidate evolved mechanisms of metabolic overproduction and dependence and to identify the extent to which the genetic and metabolic architectures imposed constraints on this process.

Importantly, evolved communities exhibited complex, multifaceted, and non-trivial metabolic interactions that were not necessarily optimized at the community level; useful metabolites were often excreted by one species but not utilized by its partner and other metabolites were cross fed without evolving complete dependency. Such “messy” interactions and missed metabolic opportunities are a reasonable outcome of selfish species evolution in the absence of explicit selection for interaction, and could also occur in natural communities. Another potential contributor to this complexity is the interconnectedness of different metabolic phenotypes induced by the genetic and metabolic architecture. Indeed, our analysis has demonstrated that genes were often associated with the excretion and/or production of multiple different metabolites. This interconnectedness may also account for the relatively frequent occurrence of provider species utilizing metabolites excreted by their dependent partners, where the gene retention and loss events that cause a dependent relationship in one direction may also facilitate emergence of a reciprocal cross-feeding relationship. Interestingly, however, in our simulations, metabolic *dependency* usually involved a single metabolite, while real world mutualistic endosymbionts often exchange and are dependent on multiple metabolites [13]. One potential explanation is that in our model bacteria continue to grow optimally (given the metabolic capacities encoded by their reduced genome), whereas in reality extreme genome reduction likely impacts cell regulation and control of growth, potentially causing cells to excrete a larger variety of useful metabolites that could be beneficial to their partners. Our analysis also suggests that the likelihood of missed metabolic opportunities may vary across metabolites, with some metabolites (e.g., cytidine, succinate, and acetate) being excreted as relatively similar frequencies in all interaction types and others (e.g., serine and thymidine) being rarely excreted in independent communities.

Our analysis additionally demonstrated how functional diversification leads to metabolic cooperation, where each species retains certain metabolic capacities that the other species has lost. Given a diversification process, it is interesting, however, to speculate about what causes one community to evolve a commensal interaction and another to evolve a mutualistic interaction. We found, for example, that provider species had on average a slightly larger genome than independent species, suggesting that a provider state is the outcome of more constrained evolutionary trajectories that end with larger minimal genomes. Our finding that dependent species in fact tend to excrete more metabolites at the beginning of the evolutionary process might further imply that early ‘wasteful’ behavior may contribute to the evolution of dependence. Another interesting outcome of our results was the dichotomy observed when a new metabolite became available, with species either starting to consume it immediately and later becoming dependent on it or never consuming it at all. These missed opportunities seem to be examples where the evolutionary events that occurred before the availability of the metabolite precluded utilization of that metabolite by potentially losing the necessary transporter or other reactions necessary for uptake. With this in mind, the non-essential cross feeding observed in commensal communities may simply represent communities that were on the path toward mutualism, but where cross-feeding emerged too late in the evolutionary process when the providers have already lost genes that would be necessary for dependence.

Despite these exciting results, there are clearly some caveats in the framework used in this work. For example, our framework assumes that bacteria grow selfishly, and accordingly cross-feeding often requires extensive gene deletions to force excretion of useful metabolites. In reality bacteria can be leaky and release metabolites into their environment even without mutations [17,24]. Another drawback stems from the fact that FBA does not take into account factors such as entropy or pH. For example, the emergence of formate cross-feeding that occurred in our simulations might be less biologically feasible because excess formate accumulation inhibits *E. coli* growth and acidifies the local environment [44]. Additionally, we model genome reduction as occurring one gene at a time [45], but do not account for the possibility of simultaneous loss of larger genomic regions [46]. Such a process could give rise to different patterns, although a study of a single reductively evolving species that examined both evolutionary regimes did not observe qualitative differences [36]. Moreover, in consideration of simulation time, in this study we only considered two-genotype communities. It would be interesting to expand our framework and to model the evolution of more complex communities or to account for spatial heterogeneity [47,48]. Finally, two decisions in the design of our study that likely had significant impacts on the outcomes were the media and the fitness cutoff. The media used is M9 minimal media, which was chosen to be more permissive of cross-feeding than rich media. Clearly, choosing different media or different limiting concentrations could impact the type of interactions that would evolve [49]. The fitness cutoff used was chosen as an intermediate value between the two cutoffs used by [36], and as shown in the supplementary text was found to have a significant effect on the frequency of evolved commensalism and mutualism. Since this fitness cutoff represents the strength of selection, a permissive fitness cutoff (as the one used in our study) allows genetic drift to play a dominant role in determining evolutionary trajectories, in agreement with the balance between selection and genetic drift hypothesize to govern the evolution of endosymbionts [26].

Looking forward, the framework presented in this study could be broadly relevant for improving our understanding of how mutualistic relationships can naturally emerge between bacterial species. This, in turn, would facilitate a deeper understanding of both simple communities, as in the case of insect endosymbionts, and significantly more complex communities, such as those inhabiting the human gut. Moreover, translationally, our approach could be useful to aid and inform the design of dependencies between bacterial species in order to increase the stability and reliability of synthetically constructed bacterial communities or interventions.

## Methods

### Evolution Simulation

The evolution simulation was initiated with a pair of genome scale metabolic models. For this study all simulations were initiated with two identical copies of the iAF1260 *E. coli* model [50]. This model includes 1260 genes, 2382 reactions, and 1668 metabolites (which includes extracellular, periplasmic, and cytoplasmic versions of some of the same metabolites). 304 of those metabolites can be exchanged with the external environment. During each step in the evolutionary process, a gene in one of the two species was chosen uniformly at random from the set of all genes still retained by the two species. The chosen gene and all the metabolic reactions that depend on this gene were deleted from the species’ model. The fitness effect of this deletion in the context of the community was determined using the co-culture growth model (see below) to evaluate the growth rate of the reduced model when grown with the current model of the partner species. If the calculated fitness effect (when compared to the fitness of that species prior to this gene deletion) was positive, neutral, or smaller than the chosen cutoff (cutoffs used include 1%, 5%, and 10%), the deletion became permanent and the process repeated with the reduced model. However, if the fitness effect exceeded the cutoff, the deletion was considered to be too harmful to occur and the process repeated until a gene that could be deleted was found. This evolutionary process continued until deletion of any remaining gene from either of the two species would cause a drop in fitness exceeding the cutoff, in which case the simulation ended. The simulation also ended if the chosen gene deletion in one species (i.e., a gene deletion that was relatively harmless for that species) caused the other species to drop significantly in fitness (>50%) in the co-culture. Such simulations, where a partner species was no longer being supported, represent collapsed communities.

### Co-Culture Growth Simulation

The co-culture growth simulation was based on a previously introduced dynamic Flux-Balance Analysis framework and is described in more detail elsewhere [37]. Briefly, given a multi-species community inhabiting a shared medium, the framework assumed that at each time step, each species grew optimally given the current concentration of metabolites in the medium (i.e., selfish growth), and then updated the abundance of each species and the concentration of metabolites in the medium based on the predicted growth and activity of each species. Specifically, at each time step, the framework first calculated the upper bound on metabolites’ uptake for each species based on the concentration of metabolites in the medium and the cell density of each species. A Flux Balance Analysis (FBA) was then used to determine the fluxes through each species’ reactions given these uptake constraints by maximizing the species’ biomass production (as a proxy for growth). A second optimization was performed to minimize the total flux through all reactions while keeping the biomass production fixed at the maximum rate (representing a minimization of enzyme usage). The predicted growth rate of each species and the predicted rates at which each species uptakes and excretes various metabolites were then used to update the cell density and concentrations of metabolites in the medium. The process was then repeated at the next time step.

For the purpose of this study, each co-culture simulation consisted of 8 steps of 0.125 hours followed by 4 steps of 0.5 hours. This provided a more accurate account of species growth at the initiation of any potential interaction, while still providing information about the co-culture growth at a longer time scale. The growth rates at the last time point (i.e., after 2.5 hours) were used as a measure of each species’ fitness. Both species started at a biomass of 0.01 grams dry mass in 1L volume for mono-culture or 2L for co-culture, resulting in the same cell density for both (which is equal to about 4*10^7 cells per liter for *E. coli*). The species were grown on a medium based on M9 minimal media [51], containing sodium, chloride, sulfate, inorganic phosphate, potassium, magnesium, ammonia, glucose, water, hydrogen, and oxygen. In addition the metals copper, iron, molybdate, manganese, zinc, nickel, and cobalt were included as they are necessary for growth of the *E. coli* model. These metabolites were all present in the medium at an excess concentration of 10M to ensure exponential growth for the entire course of the co-culture simulation. A low concentration (0.0001 mM) of jumpstart metabolites were also included to allow growth of obligate mutualistic pairs (see below). FBA solutions were calculated using glpkmex, a Matlab interface for GLPK, GNU Linear Programing Kit. GLPK version 4.54 was used, and glpkmex version 2.11.

### Jumpstarting Mutualistic Growth

Simulating the growth of species that evolved to be obligate mutualists with a dynamic FBA model has the inherent problem that neither species is able to grow initially on the minimal medium (and consequently will not excrete any of the byproducts needed to allow the other species to grow). In biological systems this problem can be overcome by heterogeneity in the growth phenotypes of individual cells, nutrients released by dead cells, or trace nutrients present in the environment. Rather than simulating diverse growth phenotypes or cell death, in this study we jumpstarted mutualistic growth by supplementing the minimal growth medium described above with trace amounts of potentially necessary metabolites. The set of these “jumpstart metabolites” was determined by identifying metabolites that could be produced by non-transfer reactions still present in at least one of the two species (even if the pathway was not complete). This set therefore represented an upper bound on which metabolites could be exchanged. Jumpstart metabolites were initialized at a low concentration of 0.0001 mM. To ensure that species that utilized these metabolites for growth could eventually be supported by the production of these metabolites by the partner species (rather than continually relying on the trace amounts of these metabolite provided at the beginning of the simulation), at 1 hour into the growth simulation, this same low concentration (0.0001 mM) was subtracted from each jumpstart metabolite.

### Filtering Completed Simulation Runs

Simulation runtime considerations necessitated using a relatively limited time resolution in the co-culture growth simulation (see above). To confirm that the evolved communities were not affected by this, for each completed simulation we ran additional co-culture growth simulations on the resulting minimal models using a finer time resolution. Specifically, co-culture growth was simulated until the medium was exhausted with time steps of 0.1 hours, using otherwise the same conditions as the co-culture growth model employed during evolution (including removing the jumpstart metabolites at 1 hour). Community growth was deemed to have been accurately simulated if:

1. Glucose eventually ran out, indicating that the two species were able to continue growing stably.
2. Both species were able to continue growth until this exhaustion of the media. Growth of both species must have been at least 50% of their measured fitness value within the last hour before all growth ended.
3. The growth rate of both species at 2.5 ± 0.2 hours was at least 90% of their fitness value as measured during the course of the evolution.

Simulations that failed any of these three criteria were excluded from the downstream analysis. Of the 16377 completed simulation runs, 16, 317 (99.6%) passed this filtering step.

### Determining Interaction Type

Interaction type was determined by comparing the fitness of each species when grown in co-culture with its fitness when grown in mono-culture. If the fitness of a species at a given time point was zero in mono-culture and non-zero in co-culture, the species was labeled as dependent at that time. If it had non-zero growth in both mono-and co-culture it was labeled as independent. Communities were labeled by the relationships between the two species: If both species were independent, the community was labeled as independent. If one species was dependent and the other was independent, the community was labeled as commensal. If both species in a community were dependent, the community was labeled as mutualistic. Within commensal communities, the dependent species was referred to as ‘dependent’ and the independent species was referred to as ‘provider’.

### Determining Metabolic Dependencies

For each evolved species we determined what metabolites it depends on (if any). To this end, we first identified metabolites that were being exchanged between the two species at the final co-culture growth time point by finding exchange reactions for which the two species had fluxes of opposite sign. The growth of each dependent species in the pair was then assayed on minimal medium supplemented with all possible combinations of these exchanged metabolites, using a single time step mono-culture growth model, to identify the smallest set of supplement metabolites that allowed it to grow at >50% its growth rate in co-culture. If no combination of supplement metabolites allowed such growth the search was expanded to include all combinations of metabolites present in the medium at the end of the co-culture simulation and that were not part of the minimal media (such metabolites could have been excreted by the provider at previous time steps).

### Pathway Analyses

Pathways annotations for each gene in the model were obtained from the Kyoto Encyclopedia of Genes and Genomes (KEGG) [52]. To identify enrichment of KEGG pathways in subsets of genes (e.g., those that were deleted at significantly different rates between interaction types), we generated 100,000 random subsets of genes of the same size and compared the total number of genes associated with each pathway in the real set to the number of genes associated with that pathway in random sets.

### Measuring Genome Similarity

To compare the similarity of two genomes (e.g., in an evolved community), we used the Jaccard similarity coefficient. We then used a two-sample t-test to test for significant differences in similarity between different types of communities.

### Identifying Co-Retained Gene Pairs

To identify gene-gene co-occurrence relationships, we examined all pairwise combinations of genes that were both retained and deleted at least three times. Since many genes perfectly co-varied with each other across simulations, we first grouped genes into sets of perfectly co-varying genes and identified co-occurrence relationships between these sets. For each pair of gene sets, we found the number of species that had retained each set and the number of species that had retained both sets and used a hypergeometric test to determine whether these sets have been co-retrained significantly more or less often than expected by chance (at 1% false discovery rate; [53]). Test for enrichment of shared pathways among the significant gene set pairs was done by permuting the connections between pairs.

### Identifying Significant Gene Deletion Ordering

Given a pair of genes, A and B, we recorded the number of times gene A in the dependent was deleted before gene B in the provider. We then used a permutation-based assay, permuting the time of deletion (measured as the position in the ordering of all gene deletions in that simulation) of each gene between all providers or dependents from commensal communities in which that gene had been deleted. Gaps and overlaps in the resulting permuted gene deletion histories were resolved by shifting deletions into gaps and randomly breaking ties. The number of times gene A in the dependent was deleted before gene B in the provider in the original data was compared to this number in the permuted data to identify significantly common ordered pairs of gene deletes (at 1% false discovery rate).

### Gene-Metabolite Connections

To identify correlations between retention or deletion of specific genes and metabolic phenotypes, we considered all genes that were both deleted and retained at least 10 times and all metabolites that were depended upon at least 10 times. For every pair of such genes and metabolites, we compared the frequency of deletion of that gene in commensal species that are dependent on that metabolite to the frequency of deletion of that gene in independent species. This was repeated for commensal species that provided the metabolite their partner depends upon, and in both cases instances of genes being deleted more often or retained more often in species with that metabolic phenotype were identified (at 1% false discovery rate).

